# Epistasis and Entropy

**DOI:** 10.1101/014829

**Authors:** Kristina Crona

## Abstract

Epistasis is a key concept in the theory of adaptation. Indicators of epistasis are of interest for large system where systematic fitness measurements may not be possible. Some recent approaches depend on information theory. We show that considering shared entropy for pairs of loci can be misleading. The reason is that shared entropy does not imply epistasis for the pair. This observation holds true also in the absence of higher order epistasis. We discuss a refined approach for identifying pairwise interactions using entropy.

## 1. INTRODUCTION

Epistasis tends to be prevalent for antimicrobial drug resistance mutations. Sign epistasis means that the sign of the effect of a mutation, whether good or bad, depends on background Weinreich et al. (2005). Sign epistasis may be important for treatment strategies, both for antibiotic resistance and HIV drug resistance Goulart et al., 2013; Desper et al., 1999; Beerenwinkel et al., 2007 a). For instance, there are sometimes constraints on the order in which resistance mutations occur. A particular resistance mutation may only be selected for in the presence of another resistance mutation. It is important to identify such constraints. A first question is how one can identify pairwise epistasis in a large system. We will discuss entropy (Shannon, 1948) and epistasis. Information theory has been used for HIV drug resistance mutations (Gupta and Adami, 2015) and more extensively for analyzing human genetic disease (e.g. Dong et al., 2008; Kang et al., 2008; Streiloff et al., 2010). For recent review articles on epistasis and fitness landscapes see e.g. Hartl (2014); Kondrashov and Kondrashov (2014), and for an empirical perspective (Szendro et al., 2012).

## 2. RESULTS

It is well established that genotypes are expected to be in equilibrium proportions if there is no epistasis in the system, i.e., if fitness is multiplicative. For instance, if two rare mutations have frequencies *p* and *q*, then the frequency of the genotype combining the two mutations is expected to be close to *pq*. This statement holds true regardless if recombination occurs or not (Otto and Lenormand, 2002).

We will explore the relation between entropy and epistasis for a system with constraints as described in the introduction.

Consider a 3-locus balletic system where a mutation at the first locus confers resistance, whereas mutations at the second and third loci are only selected for in the presence of the first mutation (otherwise they are deleterious). We represent the case with a fitness graph (Crona et al., 2013) (Figure 1). As conventional, 000 denotes the wild-type. For instance, one obtains a system with the fitness graph as in Figure 1 for the log-fitness values

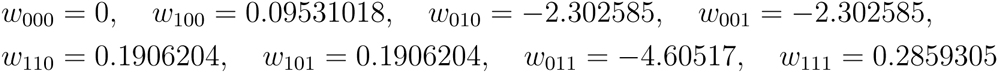

**Figure 1.**
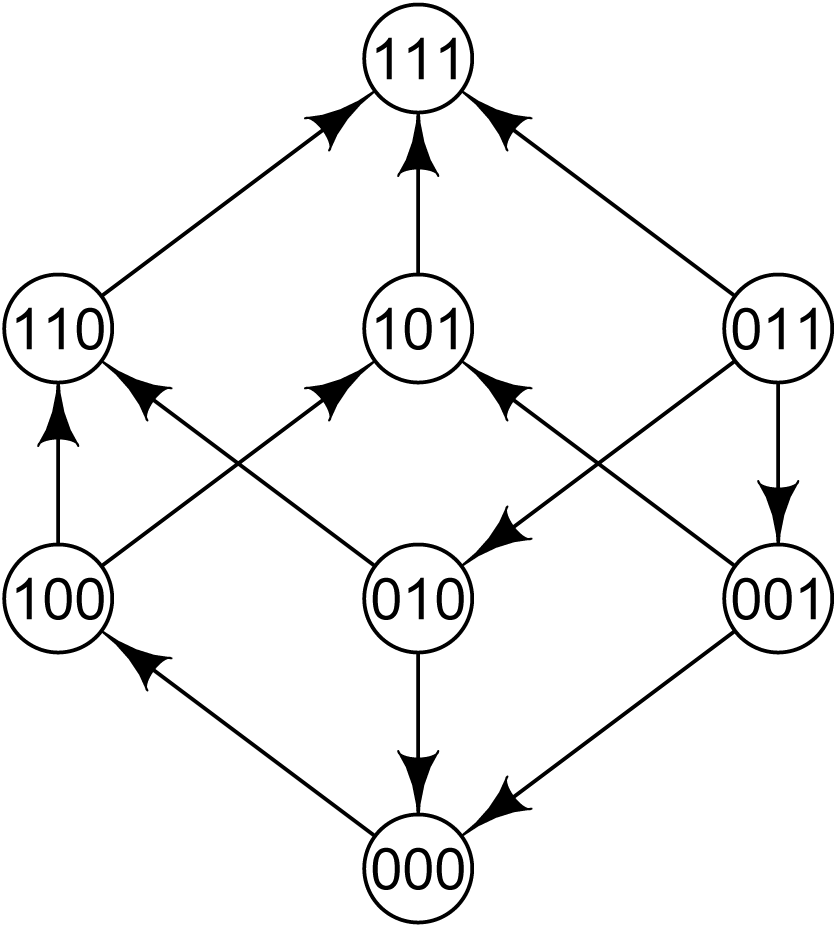
systems

The gene interactions for a 3-loci system can be described by the sign pattern of 20 circuits, or minimal dependence relations (Beerenwinkel et al., 2007 b). The relevant two-way interactions in this context be described by the six circuits corresponding to the faces of the 3-cube. Specifically,

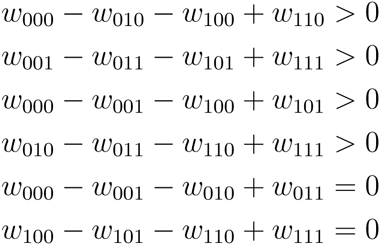

The four inequalities express that there is positive epistasis for the first and second loci, as well as for the first and third loci. The two equalities show that there is no epistasis for the second and third loci, regardless of background. The total 3-way epistasis is zero as well,

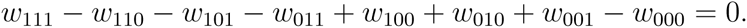

Higher order gene interactions have also been described using Walsh coefficients (Weinreich et al., 2013; Poelwijk et al, 2015). For this landscape the Walsh coefficient *E*_011_ = 0, which indicates an absence of background averaged epistasis for the second and third loci.

We will consider entropy during the process of adaptation for this landscape. The starting point for adaptation is the wild-type 000. We use a standard Wright-Fisher model for an infinite population with mutation rate *μ* = 10^−7^. The gene frequencies and shared entropy after the given number of generations are listed in the table.

**Table 1.** Gene frequencies and shared entropy *I*(2, 3) for an infinite population with mutation rate 10^−7^.

| generations | 000 | 100 | 010 | 001 | 110 | 101 | 011 | 111 | $I(2,3)$ |
| --- | --- | --- | --- | --- | --- | --- | --- | --- | --- |
| 130 | 0.7692 | 0.1850 | 0 | 0 | 0.0214 | 0.0214 | 0 | 0.0031 | 0.003206041 |
| 140 | 0.4834 | 0.3015 | 0 | 0 | 0.0904 | 0.0904 | 0 | 0.0343 | 0.01736237 |
| 146 | 0.2723 | 0.3008 | 0 | 0 | 0.1597 | 0.1597 | 0 | 0.1075 | 0.02335234 |
| 150 | 0.1569 | 0.2539 | 0 | 0 | 0.1974 | 0.1974 | 0 | 0.1944 | 0.0211462 |
| 160 | 0.0229 | 0.0959 | 0 | 0 | 0.1934 | 0.1934 | 0 | 0.4943 | 0.006950302 |
| 170 | 0.0020 | 0.0216 | 0 | 0 | 0.1132 | 0.1132 | 0 | 0.7501 | 0.001270666 |

The shared entropy for the second and third loci differs from zero. However, there is no 2-way epistasis for the pair of loci.

By extrapolation, consider an analogous system for *L*-loci. Then *L* − 1 mutations are selected for only if the first mutation has occurred, but there are no other interactions. One would get non-zero shared entropy for 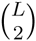 pairs of loci, although there is 2-way epistasis for *L* − 1 pairs of loci only.

### 2.1. A pair with no epistasis and maximal shared entropy

The landscape

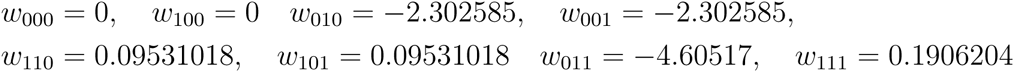
is closely related to the previous example. Indeed, the two-way interactions can be described by the sign pattern

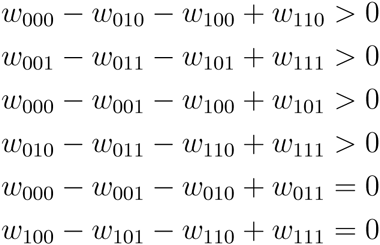
and the total 3-way epistasis is zero:

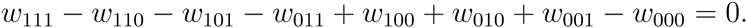

Also in this case, there is no epistasis for the second and third loci. Mutations at the second and third loci are selected for only in the presence of a mutation at the first locus. However, this fitness landscape differs from the previous example in that a mutation at the first locus is neutral for the wild-type.

Suppose that 50 percent of hosts start a new treatment with 000 viruses, and 50 percent start with the 100 genotype. That could be realistic, for instance if the 100 genotype had some resistance to a previously used drug. By assumption, eventually one would have about 50 percent 000 genotypes and 50 percent 111 genotype in the total population. Then *I*(2, 3) = 2 although there is no epistasis for the second and third loci. This example also points at a fundamental problem relating pairwise epistasis and entropy. At the time when we have 50 percent 000 genotypes and 50 percent 111 genotypes, obviously no method can reveal pairwise epistasis.

### 2.2. A refined approach

We will discuss a refined approach for identifying pairwise epistasis. Suppose that we have identified shared entropy for a particular pair of loci {*k*, *l*}. Let 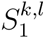 denote the set of loci such that the shared entropy

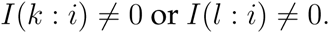

Let 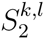 denote the set of loci with non-zero shared entropy for some locus in S_1_, and so forth. Let 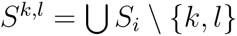.

Let *v* denote one of the 2^|*S*|^ possible states for *S*, and consider the subsystem of genotypes determined by *v*. If the shared entropy *I^v^*(*k*: *l*) = 0 for all *v*, then there is no indication of of epistasis for {*l*, *k*}.

We can apply the refined approach for the second and third loci in our example where *I*(2, 3) = 2. Then

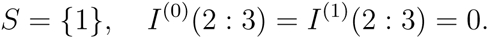

Consequently, there is no indication of epistasis for the second and third loci.

The described method could be useful for identifying cases with shared entropy and no epistasis. However, it remains to explore to what extent the method is useful in a more general setting.

## 3. DISCUSSION

We have demonstrated that shared entropy for two loci does not imply epistasis for the pair. This observation holds true also in the absence of 3-way epistasis in a single environment. Entropy based approaches to epistasis are coarse. We have discussed a refined approach which filters out some cases where shared entropy depends on states at other loci.

There are obviously other reasons for caution in interpretations of entropy for drug resistance mutations. Different drugs constitute different environments. Some resistance mutations may be correlated if they are beneficial in the presence of a particular drug, but not for other drugs. In such cases entropy would not not imply epistasis.

Our results show that observations on entropy and epistasis based on 2-locus systems can be misleading for general systems. From a theoretical point of view, a better understanding of large systems would be useful for handling drug resistance data.

## 4. METHODS

Let *x* and *y* be discrete random variables with states *x*_1_, *x*_2_ and *y*_1_, *y*_2_. Let *p*_*i*_ denote the frequency of *x*_*i*_, and *p*_*ij*_ the frequency for the combination of *x*_*i*_ and *y*_*j*_. The entropy (Shannon, 1948) *H*(*x*) and the joint entropy *H*(*x*, *y*) are defined as

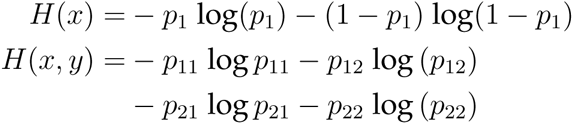

The shared entropy is the quantity *I*(*x* : *y*) = *H*(*x*) + *H*(*y*) *− H*(*x*, *y*).

In general *I* (*x* : *y*) ≥ 0, and the shared entropy is a measure of dependence.

